# Ionophore antibiotic X-206 is a potent and selective inhibitor of SARS-CoV-2 infection *in vitro*

**DOI:** 10.1101/2020.06.14.149153

**Authors:** Esben B. Svenningsen, Jacob Thyrsted, Julia Blay-Cadanet, Han Liu, Shaoquan Lin, Jaime Moyano Villameriel, David Olagnier, Manja Idorn, Søren R. Paludan, Christian K. Holm, Thomas B. Poulsen

**Affiliations:** Department of Chemistry, Aarhus University, Langelandsgade 140, DK-8000, Aarhus C, Denmark; Department of biomedicine, Aarhus University, Høegh-Guldbergs Gade 10, 8000 Aarhus C, Denmark

**Author notes:** These authors contributed equally (Esben B. Svenningsen, Jacob Thyrsted, Julia Blay-Cadanet). Thomas B. Poulsen, Christian K. Holm **Email:**.

**Keywords:** SARS-CoV-2, coronavirus, antiviral agent, polyether ionophore, X-206

## Abstract

Pandemic spread of emerging human pathogenic viruses such as the current SARS-CoV-2, poses both an immediate and future challenge to human health and society. Currently, effective treatment of infection with SARS-CoV-2 is limited and broad spectrum antiviral therapies to meet other emerging pandemics are absent leaving the World population largely unprotected. Here, we have identified distinct members of the family of polyether ionophore antibiotics with potent ability to inhibit SARS-CoV-2 replication and cytopathogenicity in cells. Several compounds from this class displayed more than 100-fold selectivity between viral-induced cytopathogenicity and inhibition of cell viability, however the compound X-206 displayed >500-fold selectivity and was furthermore able to inhibit viral replication even at sub-nM levels. The antiviral mechanism of the polyether ionophores is currently not understood in detail. We demonstrate, through unbiased bioactivity profiling, that their effects on the host cells differ from those of cationic amphiphiles such as hydroxychloroquine. Collectively, our data suggest that polyether ionophore antibiotics should be subject to further investigations as potential broad-spectrum antiviral agents.

## Introduction

The societal impact of the novel corona virus, SARS-CoV-2, that emerged in the end of 2019,^1,2^ continues to increase and the global death toll associated with the resulting respiratory disease, COVID19, is now >400.000. The lack of effective antiviral therapies and long hospitalizations required for patients with severe COVID19 threatens to overwhelm health care systems, which will continue to be risk until an effective vaccine is developed or population immunity achieved. Given the long timeframes and uncertainties involved in both of these scenarios attempts to repurpose compounds with available human safety data are of high priority. Likewise, compounds, for which animal data is available, are of strong interest for accelerated evaluation and development, in particular compounds that display broad-spectrum antiviral activity or unusual modes-of-action.^3^

The polyether ionophores is a family of natural products with diverse biological effects.^4,5^ The compounds are most known for their inhibitory activities against gram-positive bacteria, including drug-resistant strains, and coccidian protozoa, which has led to the widespread use of some polyether ionophores as veterinary antibiotics.^6^ In addition, antiviral activities against several different families of both RNA and DNA viruses have also been documented, including HIV^4,7^, influenza^8^, and Zika virus^9^. Studies conducted in the 1970-80s report inhibitory activities of nine different polyether ionophores against transmissible gastroenteritis (TGE), which is a corona virus that can infect the small intestine of pigs. Some of these agents were subsequently found to cure the infection in baby pigs.^10^ In 2014, evaluation of a panel of 290 drugs and drug candidates against MERS-CoV and SARS-CoV, revealed that 9 different ‘ion channel inhibitors’, including the polyether ionophores salinomycin and monensin, could inhibit the cytopathogenic effect of MERS-CoV, but not SARS-CoV. The specific potency and selectivity of the two molecules against MERS-CoV were however not reported.^11^ The precise antiviral mechanism of the polyether ionophores is not currently known, and earlier studies indicate that the compounds can interfere with several steps in the viral replication cycle.^4,8,9^ Here, we conduct a focused investigation on a series of polyether ionophores for their ability to inhibit SARS-CoV-2 infection *in vitro*.

## Results and Discussion

We conducted an initial screen of 11 different naturally occurring polyether ionophores and one synthetic analog for their ability to rescue the cytopathogenic effect (CPE) induced by SARS-CoV-2 viral infection of Vero cells that overexpress the serine protease TMPRSS2. Viral infection produces a distinctive phenotype of high fluorescence cells with condensed chromatin that was quantified by microscopy (Figure 1A). Interestingly, all 11 natural products display inhibitory activities in this assay although the potencies vary significantly as do the selectivity indices (SI) to effects on cellular viability (Figure 1B). A subset, including narasin, salinomycin, and nanchangmycin, display >100-fold selectivity while nigericin, indanomycin, monensin, and lasalocid have slightly lower selectivities (50-100 fold). By contrast, the canonical Ca-ionophores ionomycin and calcimycin only show modest selectivity (5- and 18-fold). The synthetic polyether ionophore HL-201, despite having potent antibacterial activity,^12^ was unselective and scarcely active. We were, however, especially intrigued to also identify compounds with strongly enhanced selectivity and potency, maduramycin (EC_50_ = 64 nM, 313-fold selectivity), and, in particular, X-206 (EC_50_ = 14 nM, 586-fold selectivity).

**Figure 1.**
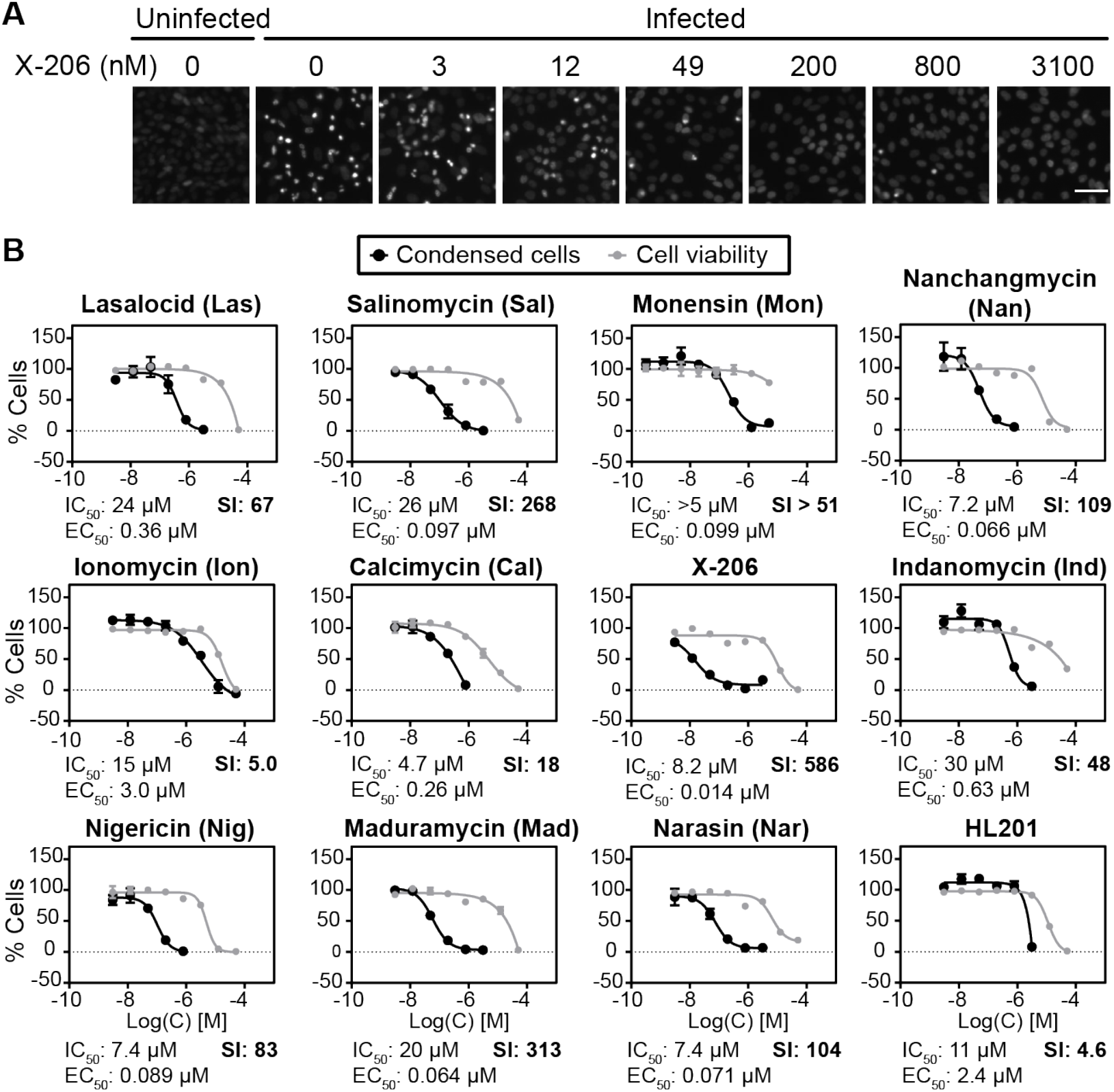
Screen of polyether ionophores for antiviral activity against SARS-CoV-2 infection in Vero-hTMPRSS2 cells. (A) Representative microscope images. Scale bar = 50 μm. (B) The relative number of viral-induced condensed cells (black lines) were counted and effects on cell viability of uninfected Vero-hTMPRSS2 cells were determined by CellTiter-Blue (grey lines). Data points are mean ± s.d (*N* = 3). SI = selectivity index.

X-206 contains a series of unusual substructures including three lactol units (Figure 2A) which, in solid-state structures, form direct interactions to bound metal ions.^13^ Interestingly, the compound was earlier found to also possess potent inhibitory activities against plasmodium parasites.^14^ To further substantiate the antiviral properties of X-206, we next assessed the inhibitory potency of the compound against viral replication in Vero-hTMPRSS2 cells using dual readouts of qRT-PCR of viral RNA and formation of the SARS-CoV-2 spike protein (Figure 2B). In this setting, X-206 displayed significant inhibition in both assays even at the lowest concentration tested (760 pM). In accord with the previous CPE-data, salinomycin also showed potent inhibition. Interestingly, hydroxychloroquine (HCQ), which was used as control, did not efficiently inhibit viral replication in Vero-hTMPRSS2 cells, but was effective in wildtype Vero cells (Figure 2C). In comparison, the potent antiviral activity of X-206 was not dependent on hTMPRSS2 expression (Figure 2B,C).

**Figure 2.**
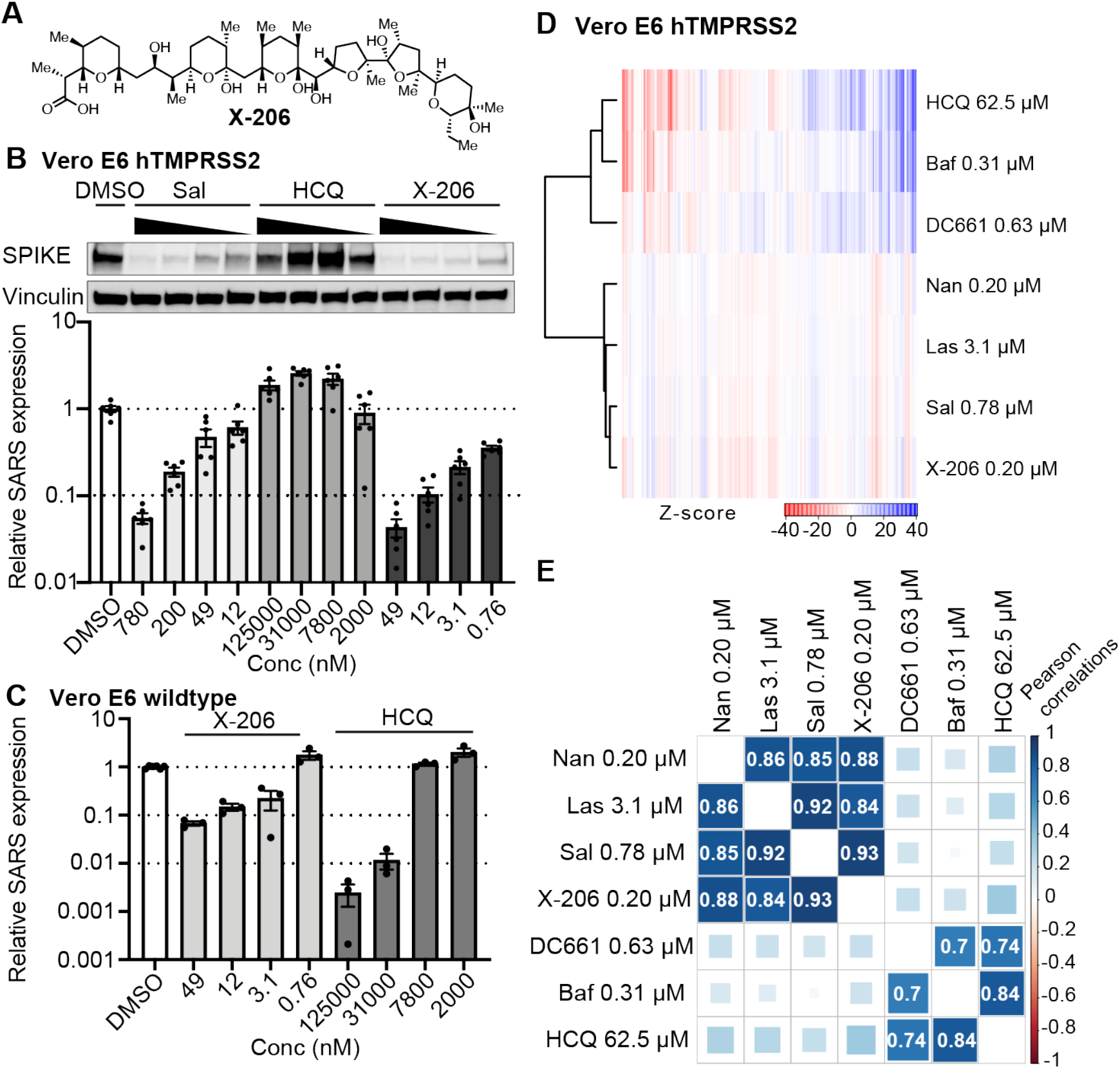
Inhibitory potency of X-206 against viral replication. (A) Molecular structure of X-206. (B) Vero E6 hTMPRSS2 cells treated for 1h followed by 24h infection. Cells were harvested for Western blot and qPCR and tested for viral titer by Spike protein or SARS-CoV-2 RNA. Western blot is representative from an experiment repeated twice with consistent outcome. qPCR is mean +/− s.d., from two distinct experiments, conducted in triplicate. (D-E) Morphological profiling of ionophores and cationic amphiphiles. (D) The Z-scored averaged bioactivity profiles. (E) Pearson correlation matrix of profiles from selected treatments ordered by hierarchical clustering.

Cationic amphiphiles, including hydroxychloroquine, are lysosomotropic agents that raise lysosomal pH and can e.g. block autophagic processes. These effects, on the host cells, are believed to be directly involved in the inhibitory activities against several types of viruses observed with many cationic amphiphiles.^15^ Polyether ionophores have also been known to accumulate in lysosomes^16^ and to inhibit autophagy^17^ and their canonical activity to facilitate exchange of metal cations for protons could also alter lysosomal pH. These similarities suggest that related mechanisms may underlie the antiviral activities observed for both cationic amphilies and polyether ionophores. We used unbiased morphological profiling^18^ to compare the effects of both types of compounds on Vero cells in the absence of viral infection (Figure 2D,E). Hydroxychloroquine, DC661, which is also a cationic amphiphile, and bafilomycin, a natural product inhibitor of V-ATPase with antiviral activity^19^, all have highly correlated (Pearson correlation coefficient *P* > 0.7) bioactivity profiles, suggesting a shared cellular mechanism (Figure 2D,E). In contrast, a selected subset of the polyether ionophores afforded bioactivity profiles that were clearly distinct from those of the cationic amphiphiles and bafilomycin, but strongly internally correlated (*P* > 0.8). The same clustering of compounds was observed in wildtype Vero cells (data not shown). The bioactive concentrations of the respective CPIs in the morphological profiling assay roughly correlated with their antiviral concentrations. This experiment suggest that polyether ionophores mediate their antiviral effects through a mechanism that is different from that of the lysosomotropic, cationic amphiphiles.

The human safety data for polyether ionophores is – to the best of our knowledge – not known and these compounds, including X-206, have toxicity in some animals. However, the fact that several members are used – safely – in the agricultural industry, and thus produced industrially, makes them relevant to consider for further studies. Our current data confirms the broad-spectrum antiviral activities of polyether ionophores, including the agricultural antibiotics salinomycin^20^, monensin and lasalocid, also against pandemic SARS-CoV-2, and furthermore reveals X-206 as a family member of exceptional potency and selectivity. Future studies will be aimed on understanding the underlying antiviral mechanism which may also shed light on the potential for further pre-clinical development.

## Materials and Methods

### Cytopathogenic Assay

20.000 Vero E6 hTMPRSS2 cells per well were seeded in 75 μL media in clear 96-well plates (Thermo Scientific, cat. no. 167008) and left to adhere overnight. Compounds (4-fold, 8-point dilution series) were diluted from 200X in DMSO to 4X in media before addition of 25 μL compound solution to the cells (final DMSO = 0.5%). Cells were treated in triplicate. After 1h treatment, all media was removed and 50 μL DMEM (with no additives) containing MOI 0,05 SARS-CoV-2 was added. After 1 hour 50 μL treatment media was added and left for 72 hours. Before harvest, cells were washed thrice with PBS and fixed with 1,5% Formaldehyde for 1 hour. Cells were then stained with Hoechst 33342n in PBS.

The plates were imaged at 8 sites in each well at 10X magnification in a Zeiss Celldiscoverer 7 microscope and exported and images were abberancies were removed. Cells with condensed chromatin were counted in CellProfiler 3.1.9. The count for the 8 sites each well was summed, all wells were subtracted the mean count from uninfected wells (10 wells), normalized to DMSO/media treated cells and fitted to a 4-parameter nonlinear regression in Prism 8.3.0 for Windows, GraphPad Software, La Jolla California USA, www.graphpad.com.

### Cytotoxicity Assay

20.000 Vero E6 hTMPRSS2 cells per well were seeded in 75 μL media in black 96-well plates (ThermoFisher Scientific, cat. no. 137103) and left to adhere overnight. Compounds (4-fold, 8-point dilution series) were diluted from 200X in DMSO to 4X in media before addition of 25 μL 4X compound solution to the cells (final DMSO = 0.5%). Cells were treated in triplicate. After 70.5 hours 20 μL CellTiter Blue (Promega, cat. no. G8080) was added to each well and incubated for 1.5 hours. The viability of the cells was assessed by measuring fluorescence in a plate reader (ex/em: 552±10/598±10, Tecan Spark 10M).

Data was subtracted background (CellTiter blue in media, no cells) normalized to DMSO/media treated cells and fitted to a 4-parameter nonlinear regression in Prism 8.3.0 for Windows, GraphPad Software, La Jolla California USA, www.graphpad.com.

### Viral replication Assay

Vero E6 cells expressing hTMPRSS2 were a kind gift of Makoto Takeda (University of Tokyo, Japan), and were cultured in DMEM (Lonza) supplemented with 10% heat inactivated fetal calf serum, 200 IU/mL penicillin, 100 μg/mL streptomycin, 600 μg/mL glutamine and 10 μg/mL blasticidin. All cell lines were regularly tested for mycoplasma contamination by sequencing from GATC Biotech (Germany). We used the SARS-CoV-2 strain #291.3 FR-4286 isolated from a patient in Germany, and kindly donated by professor Georg Kochs (Freiburg). The virus was propagated in Vero-hTMPRSS2 cells. For viral replication, 2×10^4^ Vero E6 hTMPRSS2 cells were seeded in 75 μL culture medium (w/o blasticidin). 24h after, samples were treated with appropriate compound in 25 μL DMEM for 1h. After 1h, all medium was removed and 50 μL of DMEM (with no additives) with MOI 0,05 SARS-CoV-2 was added and left for 1 hour. After 1h, 50 μL culture medium with appropriate treatment was added back and left for 72h. For WB and RT-qPCR cells were infected as described. ^3^

### Western blot analysis

Vero E6 hTMPRSS2 cells infected with SARS-Cov-2 were lysed and analyzed by Western blot as done in with Rabbit Anti-SARS CoV-2 (COVID-19) Spike (Cat No. GTX135356) (Gene Tex) and Rabbit anti-Vinculin #13901 (E1E9V XP®) (Cell signaling) as loading control. ^3^

### Reverse Transcriptase qPCR

Cells were lysed by addition of Lysis Buffer reagent (Roche) and the RNA was extracted using High Pure RNA Isolation Kit (Roche) following the manufactures protocol (Cat. No. 11-828-665-001). Validation and SARS-CoV-2 genome detection was performed with TaqMan detection systems (Applied Biosystems) based qPCR using SARS-CoV-2 specific primers and probes with the following sequences: forward primer: AAATTTTGGGGACCAGGAAC, reverse primer: TGGCACCTGTGTAGGTCAAC, probe: FAM-ATGTCGCGCATTGGCATGGA-BHQ. mRNA expressions were normalized to B-actin (Hs01060665_g1) (Thermo Fisher). ^3^

### Cell Painting assay

The experiment was conducted as described previously^21^ with the following modifications for Vero cells: 5000 cells were seeded per well, up from 4000. The following fluorophore concentrations were used: Wheat-germ agglutinin, Alexa Fluor 555 conjugate: 0.75 μg/mL; Concanavalin A, Alexa Fluor 488 conjugate: 8.75 μg/mL; Phalloidin, Alexa Flour 568 conjugate: 2.5 μL/mL; SYTO 14 Green Fluorescent Nuclei Acid Stain: 0.75 μM; Hoechst 33342: 5 μg/mL).

#### Full protocol

5000 cells were seeded into the inner 60 wells of a 96-well plate with optical bottom (Corning cat. no. 3603) in 75 μL full growth medium. After 24 h 25 μL compound solution was dosed as a 4X solution (final DMSO=0.5%). After 24 h 75 μL medium was removed and replaced with 75 μL medium containing 500 nM MitoTracker Deep Red (final C=325 nM) and plates were incubated (37 °C, 5% CO2, humid) in the dark for 30 min. Wells were then aspirated and 75 μL medium was added, before adding 25 μL 16% paraformaldehyde (Electron Microscopy Sciences 15710-S) (final PFA=4%) and incubating in the dark for 20 min. Plates were washed once with 1X HBSS (Invitrogen 14065-056) and 75 μL 0.1% (vol/vol) Triton X-100 (BDH 306324 N) in 1X HBSS was added and incubated for 15 min in the dark. Plates were washed twice with 1X HBSS before addition of 75 μL multiplex staining solution (final concentrations: Wheat-germ agglutinin, Alexa Fluor 555 conjugate: 0.75 μg/mL; Concanavalin A, Alexa Fluor 488 conjugate: 8.75 μg/mL; Phalloidin, Alexa Flour 568 conjugate: 2.5 μL/mL; SYTO 14 Green Fluorescent Nuclei Acid Stain: 0.75 μM; Hoechst 33342: 5 μg/mL) in 1% (wt/vol) BSA (Sigma-Aldrich A9647) and incubation for 30 min in the dark. Plates were then washed three times with 1X HBSS, with no final aspiration and imaged immediately in a Zeiss Celldiscoverer 7 automated microscope.

Images were analyzed using CellProfiler 2.1.1 (https://cellprofiler.org/). Per-cell profiles were averaged (mean) to per-well profiles, before Z-scoring to DMSO treated cells and averaging (mean) to per-treatment profiles in R. Profiles were compared using the Pearson correlation coefficient.

## Acknowledgments

This project has received funding from the European Research Council (ERC) under the European Union’s Horizon 2020 research and innovation programme (grant agreement No 865738). The project was also supported by Independent Research Fund Denmark (grants 9040-00117B, 0214-00001B, 9039-00078B), the Carlsbergfoundation (grant CF17-0800), Ester M og Konrad Kristian Sigurdssons Dyreværnsfond, Beckett-Fonden, Kong Christian IX og Dronning Louises Jubilæumslegat, Læge Sofus Carl Emil Friis og Hustru Olga Doris Friis’ legat, Købmand I Odense Johan og Hanne Weimann Født Seedorffs Legat, and Lundbeck foundation.

## Author Contributions

TBP, CKH, SRP conceived the study. TBP, CKH, SRP supervised the study. EBS, JTJ, CKH, and TBP designed experiments. EBS, JTJ, JBC conducted biological experiments. DO and MI contributed key reagents. HL, SL, JMV performed organic synthesis and isolation. EBS and JTJ prepared figures. TBP and CKH wrote the manuscript.

